# Neuro-Current Response Functions: A Unified Approach to MEG Source Analysis under the Continuous Stimuli Paradigm

**DOI:** 10.1101/761999

**Authors:** Proloy Das, Christian Brodbeck, Jonathan Z. Simon, Behtash Babadi

**Affiliations:** Department of Electrical and Computer Engineering, University of Maryland, College Park, MD 20742, USA; Institute for Systems Research, University of Maryland, College Park, MD 20742, USA; Department of Biology, University of Maryland, College Park, MD 20742, USA

**Author notes:** Corresponding authors (Proloy Das), (Behtash Babadi). Email addresses (Christian Brodbeck), (Jonathan Z. Simon).

**Keywords:** MEG, Temporal Response Functions, Speech Processing, Source Localization, Bayesian Estimation

## Abstract

Characterizing the neural dynamics underlying sensory processing is one of the central areas of investigation in systems and cognitive neuroscience. Neuroimaging techniques such as magnetoencephalography (MEG) and Electroencephalography (EEG) have provided significant insights into the neural processing of continuous stimuli, such as speech, thanks to their high temporal resolution. Existing work in the context of auditory processing suggests that certain features of speech, such as the acoustic envelope, can be used as reliable linear predictors of the neural response manifested in M/EEG. The corresponding linear filters are referred to as temporal response functions (TRFs). While the functional roles of specific components of the TRF are well-studied and linked to behavioral attributes such as attention, the cortical origins of the underlying neural processes are not as well understood. In this work, we address this issue by estimating a linear filter representation of cortical sources directly from neuroimaging data in the context of continuous speech processing. To this end, we introduce Neuro-Current Response Functions (NCRFs), a set of linear filters, spatially distributed throughout the cortex, that predict the cortical currents giving rise to the observed ongoing MEG (or EEG) data in response to continuous speech. NCRF estimation is cast within a Bayesian framework, which allows unification of the TRF and source estimation problems, and also facilitates the incorporation of prior information on the structural properties of the NCRFs. To generalize this analysis to M/EEG recordings which lack individual structural magnetic resonance (MR) scans, NCRFs are extended to free-orientation dipoles and a novel regularizing scheme is put forward to lessen reliance on fine-tuned coordinate co-registration. We present a fast estimation algorithm, which we refer to as the Champ-Lasso algorithm, by leveraging recent advances in optimization, and demonstrate its utility through application to simulated and experimentally recorded MEG data under auditory experiments. Our simulation studies reveal significant improvements over existing methods that typically operate in a two-stage fashion, in terms of spatial resolution, response function reconstruction, and recovering dipole orientations. The analysis of experimentally-recorded MEG data without MR scans corroborates existing findings, but also delineates the distinct cortical distribution of the underlying neural processes at high spatiotemporal resolution. In summary, we provide a principled modeling and estimation paradigm for MEG source analysis tailored to extracting the cortical origin of electrophysiological responses to continuous stimuli.

## 1. Introduction

The human brain routinely processes complex information as it unfolds over time, for example, when processing natural speech, information from lower levels has to be continuously processed to build higher level representations, from the acoustic signal to phonemes to words to sentence meaning. Quantitative characterization of the neural dynamics underlying such sensory processing is not only important in understanding brain function, but it is also crucial in the design of neural prostheses and brain-machine interface technologies.

In modeling neural activity at the meso-scale using neuroimaging modalities such as electroencephalography (EEG) and magnetoencephalography (MEG), experimental evidence suggests that linear encoding models can be beneficial in predicting the key features of sensory processing; examples include encoding models of visual and auditory stimuli (Boynton et al., 1996; Lalor et al., 2006, 2009; Ding and Simon, 2012b).

Arguably the earliest and most widely used technique to construct neural encoding models is the ‘reverse correlation’ technique, in which neural responses time-locked to multiple repetitions of simple stimuli (such as acoustic tones and visual gratings) are averaged, weighted by the instantaneous value of the preceding stimulus, to form the so-called evoked response function. Originally devised to study the tuning properties of sensory neurons (Aertsen and Johannesma, 1983; Aertsen et al., 1981; Ringach and Shapley, 2004), it was later incorporated into MEG/EEG analysis. In probing the neural response to more sophisticated stimuli such as continuous speech and video, the goal is to understand the encoding of the continuous stimuli as a whole, which is composed of both low level (e.g., acoustics) and high level (e.g., semantics) features which are bound together and distributed across time (Di Liberto et al., 2015; Brodbeck et al., 2018a).

To address this issue, techniques from linear systems theory have been successfully utilized to capture neural encoding using MEG/EEG under the continuous stimuli paradigm. In this setting, the encoding model takes the form of a linear filter which predicts the MEG/EEG response from the features of the stimulus. For example, it has been shown that the acoustic envelope of speech is a suitable predictor of the EEG response (Lalor and Foxe, 2010). These filters, or impulse response functions, play a crucial role in characterizing the temporal structure of auditory information processing in the brain, and are often referred to as Temporal Response Function (TRF) (Ding and Simon, 2012b, 2013a,b). For instance, in a competing-speaker environment in the presence of two speech streams, it has been observed that the TRF extracted from MEG response to the acoustic power consists of an early component at around 50 ms representing the acoustic power of the speech mixture, while a later peak at around 100 ms preferentially encodes the acoustic power of the attended speech stream (Ding and Simon, 2012a; Akram et al., 2016). More recent studies have expanded the TRF framework beyond the acoustic level to account for phoneme-level processing (Di Liberto et al., 2015), lexical processing (Brodbeck et al., 2018a) and semantic processing (Broderick et al., 2018).

Thanks to the grounding of the TRF model in linear system theory, several techniques from the system identification literature have been utilized for TRF estimation, such as the *normalized reverse correlation* (Theunissen et al., 2001), *ridge regression* (Machens et al., 2004), *boosting* (David et al., 2007), *and SPARLS* (Akram et al., 2017), some of which are available as software packages (Theunissen, 2007, 2010; Crosse et al., 2016; Brodbeck, 2017). While these methods have facilitated the characterization of the functional roles of various TRF components in sensory and cognitive processing of auditory stimuli, they predominantly aim at estimating TRFs over the MEG/EEG sensor space. While recent studies, using electrophysiology in animal models and ECoG in humans, have provided new insights into the cortical origins of auditory processing (see, for example, Mesgarani et al., 2008; Mesgarani and Chang, 2012; Pasley et al., 2012; Mesgarani et al., 2014), they do not account for the whole-brain distribution of the underlying sources due to their limited spatial range. As such, the whole-brain cortical origins of the TRF components are not well studied.

To address this issue using neuroimaging, current dipole fitting methods have been utilized to map the sensor space distribution of the estimated TRF components onto cortical sources (Lalor et al., 2009; Ding and Simon, 2012a). Given that the processing of sophisticated stimuli such as speech is known to be facilitated by a widely distributed cortical network, single dipole sources are unlikely to capture the underlying cortical dynamics. More recent results have used the minimum norm estimate (MNE) source localization technique to first map the MEG activity onto the cortical mantle, followed by estimating a TRF for each of the resulting cortical sources (Brodbeck et al., 2018b). While these methods have shed new light on the cortical origins of the TRF, they have several limitations that need to be addressed. First, the ill-posed nature of the MEG/EEG source localization problem under distributed source models results in cortical estimates with low spatial resolution (Schoffelen and Gross, 2009; Baillet et al., 2001). Given the recent and ongoing advances in MEG/EEG source localization towards improving the spatial resolution of the inverse solutions (see, for example, Gorodnitsky et al., 1995; Sato et al., 2004; Friston et al., 2008; Haufe et al., 2008; Wipf et al., 2010; Wipf and Nagarajan, 2009; Fukushima et al., 2012, 2015; Wu et al., 2016; Gramfort et al., 2013; Dannhauer et al., 2013; Knösche et al., 2013; Babadi et al., 2014; Krishnaswamy et al., 2017; Pirondini et al., 2018; Liu et al., 2019), it is tempting to simply use more advanced source localization techniques followed by fitting TRFs to the resulting cortical sources. However, these techniques are typically developed for the event-related potential (ERP) paradigm (Handy, 2005; Gazzaniga and Ivry, 2009; Luck, 2014) and leverage specific prior knowledge on the spatiotemporal orgranization of the underlying sources. While these assumptions bias the solution towards *source* estimates with high spatiotemporal resolution under specific repetition-based experimental settings, they do not account for the key structural properties of the underlying *neural processes* that extract information from continuous sensory stimuli. These key properties include the smoothness and/or sparsity of the response functions in the lag domain and their spatial correlation over the cortex, which may not be captured by merely enforcing spatiotemporal priors over the source domain.

Second, the single-trial nature of experiments involving continuous auditory stimuli, does not allow to leverage the time-averaging across multiple trials common in source localization of evoked responses from MEG/EEG. Third, the two-stage procedures of first fitting TRFs over the sensor space followed by localizing the peaks using dipole fitting, or first finding source estimates over the cortex followed by fitting TRFs to cortical sources, results in so-called bias propagation: the inherent biases arising from the estimation procedure in the first stage propagate to the second stage, often destructively so, and limit, sometimes severely, the statistical interpretability of the resulting cortical TRFs (see Section 3.1, for example). Finally, high resolution inverse solutions require precise forward models that are constructed based on high resolution MR scans with accurate sensor registration, which may not be readily available.

In order to address these limitations, in this paper we provide a methodology for *direct* estimation of cortical TRFs from MEG observations, taking into account their spatiotemporal structure. We refer to these cortical TRFs as neuro-current response functions (NCRFs). We construct a unified estimation framework by integrating the linear encoding and distributed forward source models, in which the NCRFs linearly process different features of the continuous stimulus and result in the observed neural responses at the sensor level. We cast the inverse problem of estimating the NCRFs as a Bayesian optimization problem where the likelihood of the recorded MEG response is directly maximized over the NCRFs, thus eliminating the need for the aforementioned two-stage procedures. In addition, to address the lack of accurate cortical surface patch statistics in the head model due to unavailability of MR scans, the NCRFs are extended to free-orientation dipoles by tripling them at each dipole location to account for vector valued current moments in 3D space. To guard against over-fitting and ensure robust recovery of such 3D NCRFs, we design a regularizer that captures the spatial sparsity and temporal smoothness of the NCRFs (e.g., minimizing the number of peaks or troughs) while eliminating any dependency on the choice of coordinate system for representing the vector valued dipole currents. While the resulting optimization problem turns out to be non-convex, we provide an efficient coordinate-descent algorithm that leverages recent advances in evidence maximization to obtain the solution in a fast and efficient manner.

We empirically evaluate the performance of the proposed NCRF estimation framework using a simulation study mimicking continuous auditory processing, which reveals that the proposed method is not only capable of identifying active sources with better spatial resolution compared to existing methods, but can also infer the orientation of the dipoles as well as the time course of the response functions accurately. Lastly, we demonstrate the utility of estimation framework by analyzing experimentally recorded MEG data from young adult individuals listening to speech for NCRFs at different hierarchical levels of speech processing. A data set, initially recorded by Presacco et al. (2016) and lacking individual MR scans, was analyzed previously by Brodbeck et al. (2018b) for source response functions using two-stage MNE followed by *boosting*-based TRF estimation. Our estimated NCRFs not only corroborate existing findings, but they are also readily interpretable in a meaningful fashion without any recourse to post-hoc processing (i.e. hierarchal clustering, sparse principal component analysis etc.) necessary for the previous study, thanks to improved spatial localization. In summary, our method successfully delineates the distinct cortical distribution of the underlying neural processes at high spatiotemporal resolution, providing new insights into the cortical dynamics of speech processing.

## 2. Theory and Methods

We develop our theory and methods for a canonical MEG auditory experiment in which the subject is listening to a single speech stream. Our goal is to determine how the different features of the speech stream are processed at different cortical stages and evoke specific neural responses that give rise to the recorded MEG data. For clarity of description and algorithm development, we first consider a single-trial experiment, and take the momentary acoustic power of the speech stream, i.e., the speech envelope, as the feature of interest. We will discuss below the more general scenarios including multiple trials, multiple speech stimuli, and multiple, and possibly competing, features reflecting different levels of cognitive processing.

### 2.1. Preliminaries and Notation

Let *e*_*t*_, 1 ≤ *t* ≤ *T* denote the speech envelope at discrete time index *t* for a duration of *T* samples taken at a sampling frequency of *f*_*s*_. We consider a distributed cortical source model composed of *M* dipole sources **d**_*m*_ = (**r**_*m*_, **j**_*m,t*_), 1 ≤ *m* ≤ *M*, where **r**_*m*_ ∈ ℝ^3^ denotes the right-anterior-superior (RAS) coordinates of the *m*^th^ dipole and **j**_*m,t*_:= [*j*_*m,t,R*_, *j*_*m,t,A*_, *j*_*m,t,S*_]^T^ ∈ ℝ^3^ denotes the dipole current vector at time *t* in the same coordinate system. The dipole locations can be obtained by standard tessellation of the 3D structural MR images of the cortex and assigning dipoles to the corresponding vertices (Baillet et al., 2001; Gramfort et al., 2014). Furthermore, the MR images can also be utilized to approximate the orientation of the current vector, assuming current flow is orthogonal to the cortical surface and replacing the dipole current vector by a scalar value (Kincses et al., 2003; Dale and Sereno, 1993). However, this approach requires precise knowledge of the cortical geometry (Haufe et al., 2008), and still might not result in ideal approximation of the cortical current orientations (Bonaiuto et al., 2019). So, it is often desirable to retain the vectorial nature of current dipoles, even though the resulting process is more complex.

Next, we assume that these current dipoles are in part stimulus-driven, i.e., each component of the current dipole relies on contributions from the preceding stimulus:

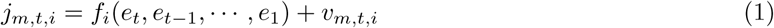

where the placeholder *i* takes the values of one of the coordinate axes, {*R, A, S*}, *f*_*i*_ is a generic function, and **v**_*m,t*_:= [*v*_*m,t,R*_, *v*_*m,t,A*_, *v*_*m,t,S*_]^T^ accounts for the stimulus-independent background activity. Following the common modeling approaches in this context (Ringach and Shapley, 2004; Lalor et al., 2006; Lalor and Foxe, 2010; Lalor et al., 2009), we take *f*_*i*_ to represent a linear finite impulse response (FIR) filter of length *L*:

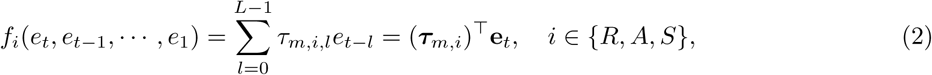

where ***τ***_*m,i*_ := [*τ*_*m,i*,0_, *τ*_*m,i*,1_ …, *τ*_*m,i,L*−1_]^T^ and **e**_*t*_ := [*e*_*t*_, *e*_*t*−1_, …, *e*_*t*−*L*+1_]^T^. Note that ***τ***_*m,i*_ can be thought of as a TRF corresponding to the activity of dipole source *m* along the coordinate axis determined by *i*. The length of the filter *L* is typically determined by a priori assumptions on the effective integration window of the underlying neural process. When stacked together, the 3D linear filters ***τ***_*m*_ := [***τ***_*m,R*_, ***τ***_*m,A*_, ***τ***_*m,S*_] ∈ ℝ^*L*×3^ are vector-valued TRFs at each source *m*, capturing the linear processing of the stimuli at the cortical level. As such, we refer to these vector-valued filters as Neuro-Current Response Functions (NCRFs) henceforth. Intuitively speaking, the 3D vector (*τ*_*m,R,l*_, *τ*_*m,A,l*_, *τ*_*m,S,l*_)^T^ is the vector-valued dipole activity at a lag of *l/f*_*s*_ second arising from a putative stimulus impulse at time 0.

Let 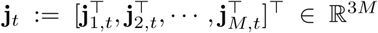 be a vector containing all the current dipoles at time *t*, and **J** := [**j**_1_, …, **j**_*T*_] ∈ ℝ^3*M* ×*T*^ be the matrix of current dipoles obtained by concatenating the instantaneous current dipoles across time *t* = 1, 2, …, *T*. Similarly, let ***V*** ∈ ℝ^3*M* ×*T*^ denote the matrix of stimulus-independent background activity, **Φ** := [***τ***_1_, ***τ***_2_, …, ***τ***_*M*_]^T^ ∈ ℝ^3*M* ×*L*^ denote the matrix of NCRFs, and **S** := [**e**_1_, **e**_2_, …, **e**_*T*_] ∈ ℝ^*L*×*T*^ denote the matrix of features. Eqs. (1) and (2) can then be compactly expressed as:

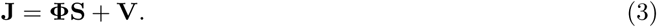

As for the sensor space, we assume a conventional MEG setting with *N* sensors placed at different positions over the scalp, recording magnetic fields/gradients as a multidimensional time series. The MEG observation at the *i*^th^ sensor at time *t* is denoted by *y*_*i,t*_, 1 ≤ *i* ≤ *N* and *t* ∈ [1, …, *T*]. Let **Y** ∈ ℝ^*N*×*T*^ be the MEG measurement matrix with the (*i, t*)^th^ element given by *y*_*i,t*_. The MEG measurement matrix is related to the matrix of current dipoles **J** according to the following forward model (Sarvas, 1987; Mosher et al., 1999; Baillet et al., 2001):

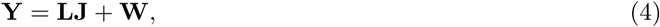

where **L** ∈ ℝ^*N*×*dM*^ maps the *source space* activity to the *sensor space* and is referred to as the *lead-field matrix*, and **W** ∈ ℝ^*N*×*T*^ is the matrix of additive measurement noise. The lead-field matrix can be estimated based on structural MRI scans by solving Maxwell’s equations under the quasi-static approximation (Hämäläinen et al., 1993).

### 2.2. Problem Formulation

Given the stimulus-driven and current-driven forward models of Eqs. (3) and (4), our main goal is to estimate the matrix **Φ**, i.e., the NCRFs. To this end, we take a Bayesian approach, which demands distributional assumptions on the various uncertainties involved, i.e., the stimulus-independent background activity and the measurement noise. For the measurement noise, we adopt the common temporally uncorrelated multivariate Gaussian assumption, i.e.,

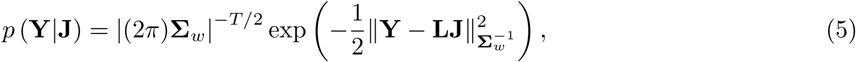

where 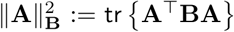 and 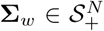 denotes the unknown noise covariance matrix. The covariance matrix **Σ**_*w*_ can be estimated from either empty-room or pre-stimulus recordings (Engemann and Gramfort, 2015). Next, let **V**_*m*_ ∈ ℝ^3×*T*^ denote the matrix of background activity at source *m*, for *m* = 1, 2, …, *M*. We adopt the following distribution for the background activity **V**:

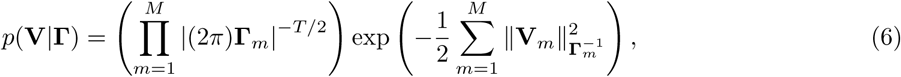

i.e., the portion of the current dipoles reflecting the background activity are modeled as zero-mean independent Gaussian random vectors with unknown 3D covariance matrix 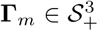. Under this assumption, Eq. (3) can be expressed as:

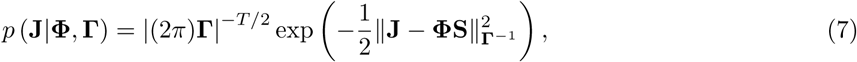

where 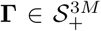 is a block-diagonal covariance matrix with its *m*^th^ diagonal block given by **Γ**_*m*_, for *m* = 1, 2, …, *M*.

Under these assumptions, the joint distribution of the MEG measurement and current dipole matrices is given by:

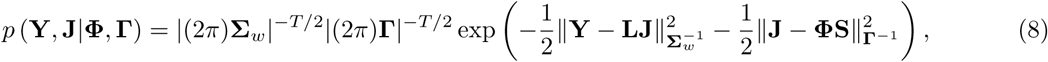

By marginalizing over **J** (see Appendix A for details), we obtain the distribution of the MEG measurement matrix parametrized by the NCRF matrix **Φ** and the source covariance matrix **Γ**:

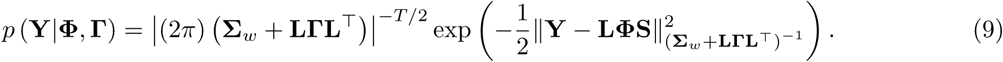

It is now possible to cast the problem of finding **Φ** as a Bayesian estimation problem, in which a loss function fully determined by the posterior distribution of NCRF matrix **Φ** given the MEG measurement matrix **Y** is minimized. In other words, if **Γ** were known, the NCRF matrix estimation would amount to the following *maximum likelihood* problem:

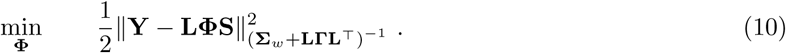

Another advantage of this Bayesian framework is the possibility of introducing regularization schemes that can mitigate the ill-posed nature of this problem, and instead work with regularized maximum likelihood problems. Note that this optimization problem makes a direct connection between the MEG measurement matrix,**Y** and the NCRF matrix **Φ** and allows us to avoid the aforementioned two-stage procedures in finding TRFs at the cortical level (Lalor et al., 2009; Brodbeck et al., 2018b).

#### 2.2.1. Regularization

As is the case in other source imaging methods, there are many fewer constraints than the free parameters determining the NCRFs. This makes the problem severely ill-posed. As such, proceeding with the maximum likelihood problem in Eq. (10) is likely to result in overfitting. In order to ensure robust recovery of a meaningful solution to this ill-posed problem, we need to include prior knowledge on the structure of the NCRFs in the form of regularization.

To this end, we construct regularizers based on a convex norm of the NCRF matrix **Φ**, to both capture the structural properties of the NCRFs and facilitate algorithm development. The structural properties of interest in this case are spatial sparsity over the cortical source space, sparsity of the peaks/troughs, smoothness in the lag domain, and rotational invariance (Ding and Simon, 2012b; Akram et al., 2017).

In order to promote smoothness in the lag domain and sparsity of the peaks/troughs, we adopt a concept from Chen et al. (2001), in which a temporally smooth time series is approximated by a small number of Gabor atoms over an over-complete dictionary 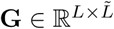, for some 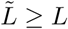 (Feichtinger and Strohmer, 2012; Akram et al., 2017). To this end, we first perform a change of variables ***τ***_*m*_ := **G*θ***_*m*_, **Φ** = **ΘG**^T^, and 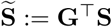, where 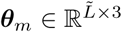 are the coefficients of the *m*^th^ NCRF over the dictionary **G** and 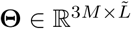 is a matrix containing ***θ***_*m*_s across its rows. Then, to enforce sparsity of the peaks/troughs, spatial sparsity, and rotational invariance, we use the following mixed-norm penalty over ***θ***_*m*_s, i.e., the Gabor coefficients:

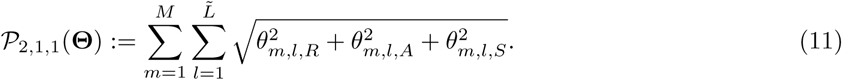

Let ***θ***_*m,l*_ ∈ ℝ^3^ be the *l*^th^ Gabor coefficient vector for the *m*^th^ NCRF. Note that the summand is ‖***θ***_*m,l*_‖_2_, which is a rotational invariant norm with respect to the choice of dipole RAS coordinate system. This structural feature allows the estimates to be robust to coordinate rotations (see Appendix B). The inner summation of ‖***θ***_*m,l*_‖_2_ (as opposed to 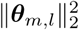) over 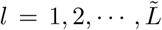 enforces group sparsity of the Gabor coefficients (i.e., the number of peaks/troughs), akin to the effect of *ℓ*_1_-norm. Finally, the outer summation over *m* = 1, 2, …, *M* promotes spatial sparsity of the NCRFs (see Fig. 1, and also Appendix B).

**Figure 1:**
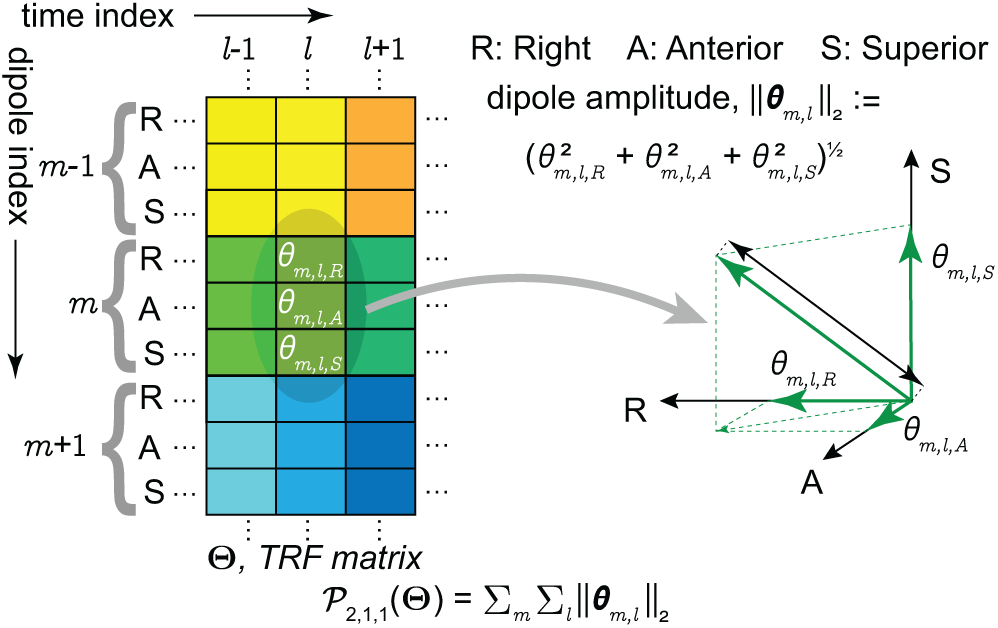
Mixed-norm penalty term 𝒫_2,1,1_(**Θ**) for regularizing the loss function. The penalty term is constructed by first isolating all 3D Gabor coefficient vectors across the dictionary elements and space, and then aggregating their *ℓ*_2_ norm. As a result, it promotes sparsity in space and Gabor coefficients, while being invariant to the orientation of the dipole currents.

Using this change of variables and regularization scheme, we can reformulate (10) as the following regularized maximum likelihood problem:

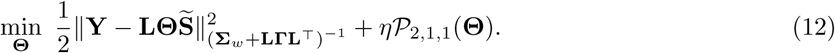

The parameter *η >* 0 controls the trade-off between data fidelity and regularization, i.e., the complexity of the resulting model grows inversely with the magnitude of *η*. This parameter can be chosen in a data-driven fashion using cross-validation (see Section 3.2). Fig. 2 provides a visual illustration of the proposed modeling and estimation paradigm.

**Figure 2:**
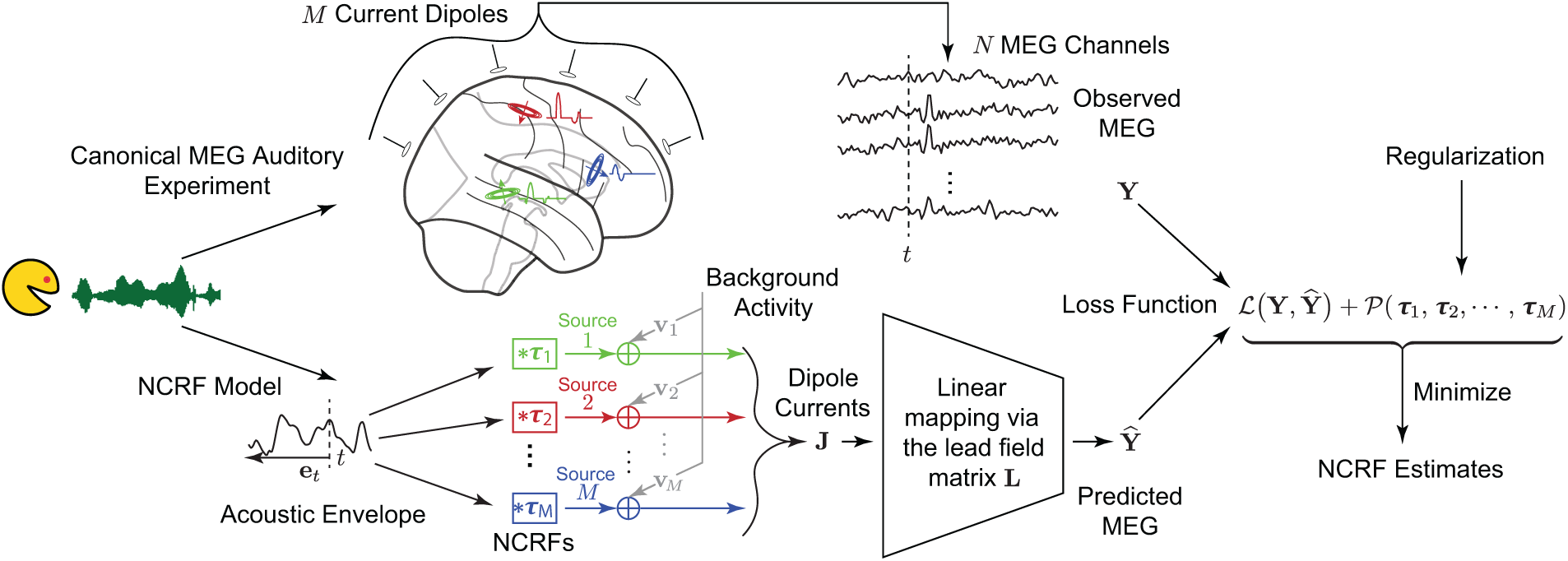
Schematic depiction of the proposed modeling and estimation framework. Upper branch: the experimental setting in which the subject is listening to speech while MEG neural responses are being recorded. Lower branch: the modeling framework in which the speech waveform is transformed into a feature variable representation, and is thereby processed via *M* linear filters (i.e., NCRFs) to generate time-varying current dipoles at each of the corresponding *M* source locations. Note that each NCRF in the lower branch corresponds to a 3D vector of dipole activity with a specific temporal profile, as shown in the upper branch with matching colors. These dipoles give rise to the predicted MEG response via a source-to-sensor mapping (i.e., the lead-field matrix). The two branches converge on the right hand side, where the NCRFs are estimated by minimizing a regularized loss function.

#### 2.2.2. Source Covariance Matrix Adaptation

Note that the objective function in Eq. (12) is convex in **Θ** and thus one can proceed to solve for **Θ** by standard convex optimization techniques. However, this requires the knowledge of the source covariance matrix **Γ**, which is unknown in general. From Eq. (7), it is evident that **Γ** implicitly offers adaptive penalization over the source space through spatial filtering. As such, the source covariance matrix serves as a surrogate for depth compensation (Lin et al., 2006), by reducing the penalization level at locations with low SNR. One data-independent approach for estimating **Γ** is based on the lead-field matrix (Haufe et al., 2008). Here, thanks to the Bayesian formulation of our problem, we take a data-driven approach to adapt the source covariance matrix to the background activity not captured by the stimulus (Stahlhut et al., 2013). One principled way to do so is to estimate both **Θ** and **Γ** from the observed MEG data by solving the following optimization problem:

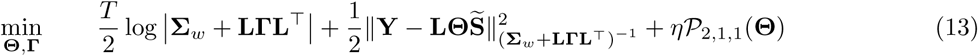

Unfortunately, the loss function in Eq. (13) is not convex in **Γ**. However, given an estimate of **Θ**, solving for the minimizer of (13) in **Γ** is a well-known problem in Bayesian estimation and is referred to as evidence maximization or empirical Bayes (Berger, 1985). Although a general solution to this problem is not straightforward to obtain, there exist several Expectation-Maximization (EM)-type algorithms, such as ReML (Friston et al., 2008), sMAP-EM (Lamus et al., 2012), and the conjugate function-based algorithm called Champagne (Wipf et al., 2010), which might be employed to estimate **Γ** given an estimate of **Θ**. In the next section, we present an efficient recursive coordinate descent-type algorithm that leverages recent advances in evidence maximization and proximal gradient methods to solve the problem of Eq. (13).

### 2.3. *Inverse Solution: The* Champ-Lasso *Algorithm*

Since simultaneous minimization of (13) with respect to both **Θ** and **Γ** is not straightforward, we instead aim to optimize the objective function by alternatingly updating **Θ** and **Γ**, keeping one fixed at a time. Suppose after the *r*^th^ iteration, the updated variable pair is given by (**Θ**^(*r*)^, **Γ**^(*r*)^), then the update rules for (*r* + 1)^th^ iteration are as given as follows:

*Updating* **Γ**

With **Θ** = **Θ**^(*r*)^, Eq. (13) reduces to the following optimization problem:

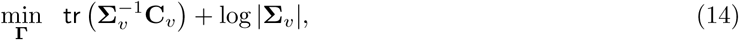

with 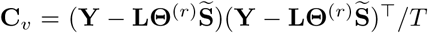 and **Σ**_*v*_ = **Σ**_*w*_ + **LΓL**^T^. Although the problem is non-convex in **Γ**, it can be solved via the Champagne algorithm (Wipf et al., 2010), which solves for **Γ** by updating a set of auxiliary variables iteratively. Though the solution **Γ**^(*r*+1)^ is not guaranteed to be a global minimum, the convergence rate is fast (with computation cost per iteration being linear in *N*), and more importantly each iteration is guaranteed not to increase the loss function in Eq. (14).

*Updating* **Θ**

Fixing **Γ** = **Γ**^(*r*+1)^, results in the following convex optimization problem over **Θ**:

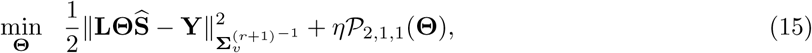

where 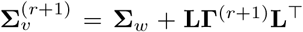. The first term in Eq. (15) is a smooth differentiable function whose gradient is straightforward to compute, and the proximal operator for the penalty term 𝒫_2,1,1_(**Θ**) has a closed-form expression and can be computed in an efficient manner (Gramfort et al., 2012). Regularized optimization problems of this nature can be efficiently solved using an instance of the forward-backward splitting (FBS) method (Beck and Teboulle, 2009; Nesterov, 2005). We use an efficient implementation of FBS similar to FASTA (Fast Adaptive Shrinkage/Thresholding Algorithm) software package (Goldstein et al., 2014) to obtain **Θ**^(*r*+1)^ from Eq. (15).

Although the loss function is not jointly-convex in (**Θ, Γ**), the foregoing update steps ensure that the loss in (13) is not increased at any iteration and stops changing when a fixed-point or limit-cycle is reached (Wright, 2015). Finally, **Γ**^0^ can be initialized according to MNE-Python recommendations for choosing the source covariance matrix in computing linear inverse operators. Also note that due to the efficiency of the overall solver, it is possible to start the optimization with several randomized initializations, and choose the best solution among several potential alternatives.

### 2.4. Extension to Multiple Feature Variables

The preceding sections focused on the case of a single stimulus feature variable, i.e., the speech envelope. However, complex auditory stimuli such as natural speech, are processed at various levels of hierarchy. Upon entering the ears, the auditory signal is decomposed into an approximate spectrogram representation at the cochlear level prior to moving further into the auditory pathway (Yang et al., 1992). Beyond these low-level acoustic features, higher-level phonemic, lexical, and semantic features of the natural speech are also processed in the brain. Thus, to obtain a complete picture of complex auditory cortical processing, it is desirable to consider response functions corresponding to more than one feature variable.

#### Algorithm 1 The Champ-Lasso Algorithm over Multiple Trials

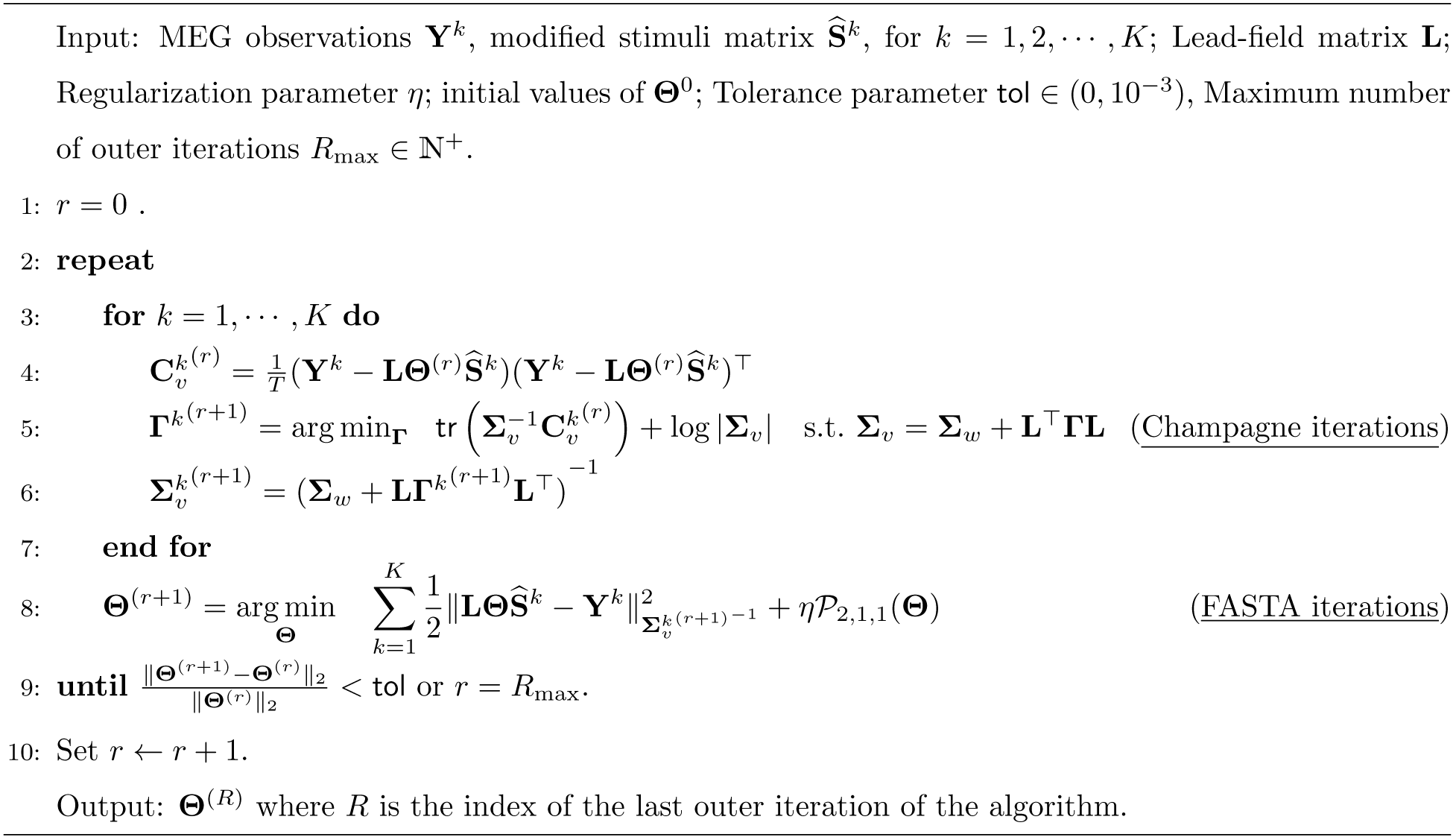

One can proceed to estimate response functions for each feature variable separately. But, since many of these features have significant temporal correlations, the resulting response functions do not readily provide unique information regarding the different levels of the processing hierarchy. To investigate simultaneous processing of these various feature variables and allow them to compete in providing independently informative encoding models, we consider a multivariate extension of the response functions (Ding and Simon, 2012b; Di Liberto et al., 2015).

Suppose that there are *F* ≥ 1 feature variables of interest. We modify Eq. (3) by replacing each column of the NCRF matrix **Φ** by *F* columns (one for each temporal response function) and each row of the stimulus matrix by *F* rows (one for each feature variable). As we will demonstrate below in Section 3, this will enable us to distinguish between different cortical regions in terms of their response latency across a hierarchy of features.

### 2.5. Extension to Multiple Trials with Different Stimuli

Next, we consider extension to *K* different trials corresponding to possibly different auditory stimuli. Let the stimuli, MEG observation, and background activity covariance matrices for the *k*^th^ trial be denoted by 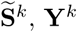, and **Γ**^*k*^, respectively, for *k* = 1, …, *K*. We can extend the optimization problem of Eq. (13) as follows:

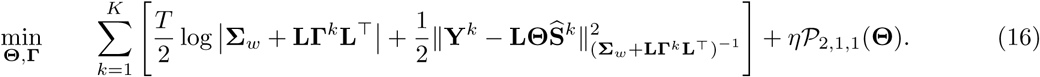

In doing so, we have assumed that the background activity is a stationary Gaussian process within a trial (with covariance **Γ**^*k*^ at trial *k*), and that the NCRFs remain unchanged across trials, which promotes integration of complementary information from different trials (without direct averaging). Note that this assumption intentionally suppresses the trial-to-trial variability of the NCRFs by adaptively weighting the contribution of each trial according to its noise level (i.e., **Γ**^*k*^), in favor of recovering NCRFs that can explain common cortical patterns of auditory processing. In contrast, if all the trials were to be concatenated or directly averaged to form a unified trial (with a single covariance matrix **Γ**), the trial-to-trial variability would not necessarily be suppressed, especially when there are few trials available. Furthermore, this formulation allows to incorporate trials with different lengths into the same framework.

The optimization problem of Eq. (16) can be solved via a slightly modified version of that presented in Section 2.3. The resulting algorithm is summarized in Algorithm 1, which we refer to as the Champ-Lasso algorithm. A Python implementation of the Champ-Lasso algorithm is archived on the open source repository Github (Das, 2019) to ease reproducibility and facilitate usage by the broader systems neuroscience community.

### 2.6. Subjects, Stimuli, and Procedures

The data used in this work are a subset of recordings presented in (Presacco et al., 2016), and is publicly available in the Digital Repository at the University of Maryland (Presacco et al., 2018). The auditory experiments were conducted under the participation of 17 young adult subjects (aged 18-27 years), recruited from the Maryland, Washington D.C. and Virginia area. The participants listened to narrated segments from the audio-book, *The Legend of Sleepy Hollow by Washington Irving* (https://librivox.org/the-legend-of-sleepy-hollow-by-washington-irving/), while undergoing MEG recording. Although the dataset contains recordings under different background noise levels, for the current work we considered recordings of two 1 min long segments of the audio-book with no background noise presented as single-speaker audio. Each of these segments was repeated three times to every individual, yielding a total 6 min of data per subject. To ensure that the participants actively engage in the listening task, they were tasked to also silently count the number of specific words that they would hear in the story.

### 2.7. Recording and Preprocessing

The data were acquired using a whole head MEG system (KIT, Nonoichi, Ishikawa, Japan) consisting of 157 axial gradiometers, at the University of Maryland Neuroimaging Center, with online low-pass filtering (200 Hz) and notch filtering (60 Hz) at a sampling rate of 1 kHz. Data were pre-processed with MNE-Python 0.18.1 (Gramfort et al., 2013, 2014). After excluding flat and noisy channels, temporal signal space separation was applied to remove extraneous artifacts (Taulu and Simola, 2006). Data were then filtered between 1 Hz to 80 Hz using a zero-phase FIR filter (using the default filter parameter options of MNE-Python 0.18.1). Independent component analysis (extended infomax, Bell and Sejnowski, 1995) was applied to remove ocular and cardiac artifacts. Finally, 60 s long data epochs corresponding to the stimuli were extracted and downsampled to 200 Hz.

### 2.8. Source Space Construction

At the beginning of each recording session, each participant’s head shape was digitized with a 3-Space Fastrak system (Polhemus), including 3 fiducial points and 5 marker positions. Five marker coils attached to these five marker positions were used to localize the head position of the participant relative to the MEG sensors. The head position measurement was recorded twice: at the beginning and end of the recording session and the average measured head positions were used. Since MR scans of the participants were not performed, the ‘fsaverage’ brain model (Fischl, 2012) was co-registered (via uniform scaling, translation and rotation) to each participant’s head, using the digitized head shapes.

A volumetric source space for the ‘fsaverage’ brain was defined on a regular grid with spacing of 7 mm between two neighboring points, and then morphed to individual participants. These morphed source spaces were then used to compute lead-field matrices by placing 3 orthogonal virtual current dipoles on each of the grid points. The computed lead-field matrices contained contribution from 3222 virtual current dipoles, after removing those within subcortical structures along the midline. No cortical patch statistics were available due to the lack of MR scans, so the current dipoles were allowed to have arbitrary orientations in 3D.

### 2.9. Stimulus Feature Variables

We included predictor variables reflecting three different hierarchical levels of speech processing, including acoustic, lexical, and semantic features. These feature variables are described in detail in (Brodbeck et al., 2018b):

- *Envelope:* The speech envelope was found by averaging the auditory spectrogram representation generated using a model of the auditory periphery (Yang et al., 1992) across frequency bands. This continuous univariate feature variable reflects the momentary acoustic power of the speech signal.
- *Word Frequency:* First, logarithmic word frequency measures, log_10_ wf, were extracted from the SUBTLEX database (Brysbaert and New, 2009) for each word. Then, a piecewise-continuous feature variable was constructed by representing each word in the speech segment by a rectangular pulse with height given by 6.33 − log_10_ wf. Note that in this coding scheme, infrequent words are assigned higher values, while common words get lower values. Windows of silence were assigned 0.
- *Semantic Composition:* Lastly, to probe semantic processing, the semantic composition patterns identified by (Westerlund et al., 2015), including adjective-noun, adverb-verb, adverb-adjective, verb-noun, preposition-noun and determiner-noun pairs, were used. To generate the feature variable, the second word in each pair was represented by a rectangular window of height 1, and 0 elsewhere. This binary-valued feature variable identifies the semantic binding of word pairs within the speech stream.

All three variables were constructed from the speech segments at the same sampling frequency as the preprocessed MEG data (i.e. 200 Hz). All feature variables were centered and scaled by their mean absolute value, to facilitate comparison of NCRF components pertaining to different feature variables.

### 2.10. Estimation Setup, Initialization and Statistical Tests

We estimated 1000 ms-long NCRFs (*L* = 200) corresponding to each of these three stimulus variables (*F* = 3). This choice leads to a high-dimensional NCRF matrix **Φ** ∈ ℝ^9666×600^. The noise covariance matrix, **Σ**_*w*_, was estimated from empty-room data using MNE-Python 0.18.1 (Gramfort et al., 2013, 2014) following an automatic model selection procedure. The regularization parameter was tuned on the basis of generalization error via a 3-fold cross-validation procedure: from a predefined set of regularization parameters (equally spaced in logarithmic scale), the one resulting in the least generalization error was chosen to estimate the NCRFs for each subject. To maintain low running time, instead of utilizing a randomized initialization scheme for **Γ**^0^, we initialized it according to the MNE-Python recommendation for source covariances. The NCRF matrix **Θ**^0^ was initialized as an all zero matrix. In the consecutive iterations of the Champagne and FASTA algorithms, a warm starting strategy was followed, i.e., initializing each iteration by the solution of the previous one.

To check whether inclusion of each of the feature variables improves the overall NCRF model significantly, the original model fit being tested for significance (i.e., its cost function evaluated at the estimated NCRF parameters) was compared against the average of three other model fits constructed by deliberately misaligning one feature variable via 4-fold cyclic permutations, using one-tailed t-tests.

To evaluate the group-level significance of the estimated NCRF components, the NCRF estimates were first smoothed with a Gaussian kernel (with standard deviation of 10 mm) over the source locations to compensate for possible head misalignments and anatomical differences across subjects. Then, at each dipole location and time index, the magnitudes of the vector-valued NCRFs were tested for significance using a permutation test via the threshold-free cluster-enhancement (TFCE) algorithm (Smith and Nichols, 2009) (see Appendix C for details).

### 2.11. Simulation Setup

Before applying the Champ-Lasso algorithm to localize NCRFs from experimentally recorded data, we assessed its performance using realistic simulation studies with known ground truth. In accordance with our experimental settings, we synthesized six 1 min long MEG data segments according to the forward model of Eqs. (3) and (4), mimicking the neural processing of the speech envelope. To this end, we simulated temporal response functions of length 500 ms (with significant M50 and M100 components) associated with dipole current sources within the auditory and motor cortices. Fig. 3A shows the simulated activity over the cortical surface at specific time lags (top and bottom panels) as well as the temporal profile of the NCRFs (middle panel). The cortical activity was simulated using patches defined over a finely-discretized source space (namely, ico-5, with average dipole spacing of 3.1 mm) with the dipole directions constrained to be normal to the ‘fsaverage’ surface patches. To make the simulation as realistic as possible, we used real MEG recordings corresponding to a different speech stream as background noise (i.e., stimulus-independent background activity), maintaining a −5 dB signal-to-noise ratio.

**Figure 3:**
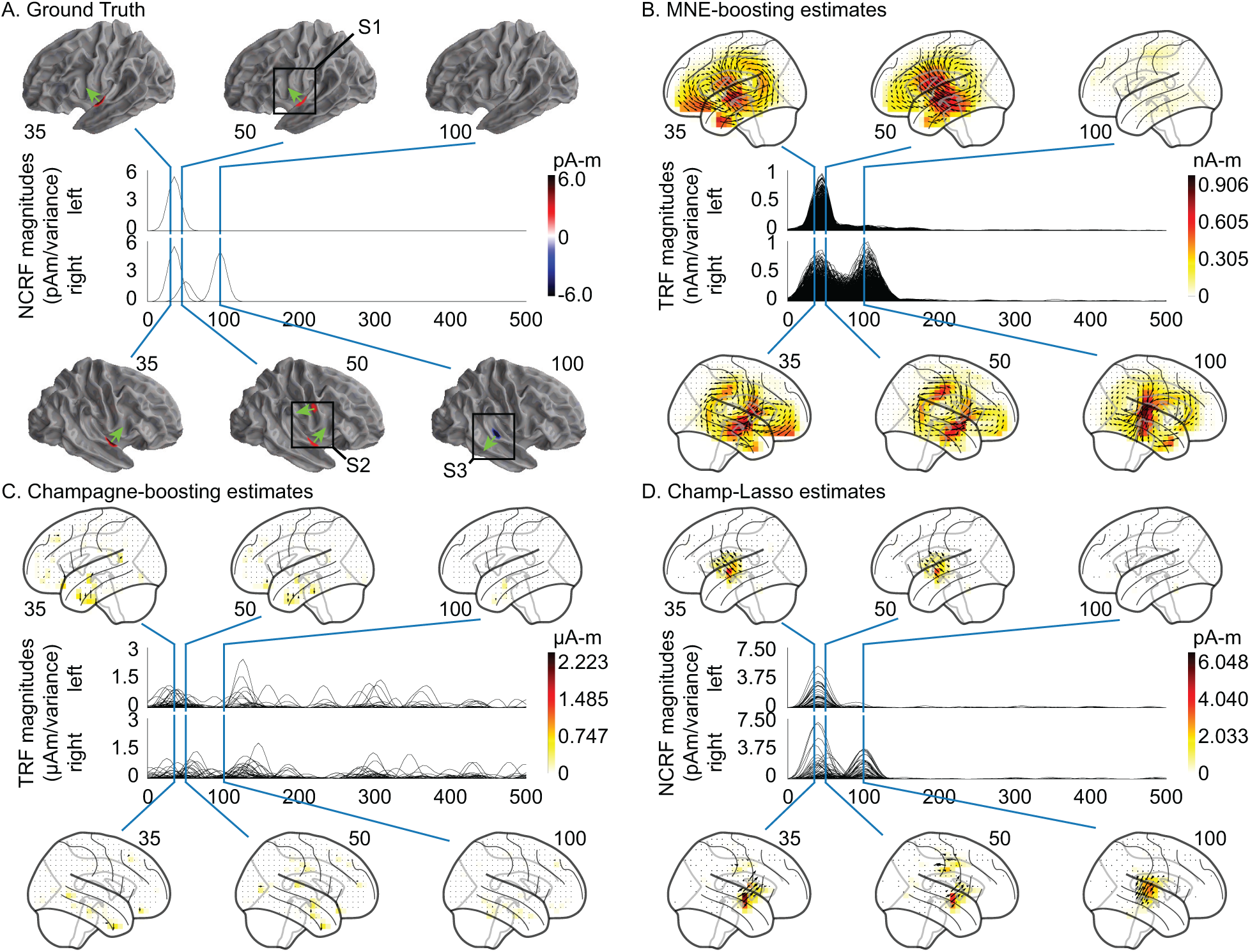
Results for a simulated auditory experiment. The top and bottom portions of each subplot pertain to the left and right hemispheres, respectively. A. The anatomical plots show the simulated neural sources normal to cortical surface, and the traces show the overlaid temporal profiles. The colorbar encodes directional intensity normal to the cortical surface (shown by the green arrows). B & C. The two-stage localized (i.e. MNE-boosting, and one of its variants, Champagne-boosting, respectively) TRFs (free-orientation) are shown on the anatomical plots, where the 3D dipoles are projected onto the lateral plane. D. NCRF estimates from the Champ-Lasso algorithm. The colorbar encodes dipole magnitudes. The spatial extent, dipole moment scale, temporal profile, and orientations of the neural sources are faithfully recovered by the Champ-Lasso algorithm, whereas the two-stage localized TRFs are either spatially dispersed (MNE-boosting) or overly sparse (Champagne-boosting) and exhibit spurious peaks in the anterior temporal and inferior frontal lobes.

In order to avoid any favorable bias in the inverse solution, we used a different source space for NCRF estimation, i.e., the aforementioned volumetric source space with unconstrained dipole orientations (Section 2.8), than the one used for simulating the data, i.e., ico-5. As a comparison benchmark, we also applied the two-stage method of Brodbeck et al. (2018b), MNE-boosting, and one of its variants, Champagne-boosting, to first localize the cortical sources using MNE and Champagne, respectively, followed by boosting with 10-fold cross-validation and *ℓ*_1_-norm error of the standardized source estimates, for independently estimating TRFs for all sources. The boosting was initialized with a zero response function, and iteratively modified it in small increments (typically 0.001) at a single time-lag in which a change led to the largest *ℓ*_1_-norm error reduction in the training set. The process stopped when the training error no longer decreased, or testing error increased in two successive steps.

In order to compare the spatial spread across different methods, we computed the *dispersion* metric as the ratio of total NCRF power outside and inside of spheres of radius *r* (for *r* = 10, 15, 20 mm) around the center of mass of the simulated cortical patches (i.e., lower is better). To quantify the response function reconstruction performance, the 3-dimensional NCRFs within radius of *r* = 15 mm around the center of mass of the simulated cortical patches were averaged and then separated into *principal orientation* and *principal time course*, using singular value decomposition. The principal orientations and time courses were compared to the ground truths using the Pearson correlation (i.e., higher is better). Finally, we quantified the *selectivity* of the principal orientation and time course in the recovered NCRFs, by the ratio of the principal singular value to the sum of all three (i.e., higher is better).

## 3. Results

### 3.1. Simulation Studies

The two two-stage localized TRFs and estimated NCRFs are shown in Fig. 3B, 3C and 3D, respectively. Since boosting tends to result in temporally sparse response functions (David et al., 2007), response functions were smoothed with a 50 ms Hamming window. The anatomical plots show the spatial response function profile at the same temporal peaks selected in Fig. 3A, with direction of the vectors projected onto the lateral plane. The Champ-Lasso algorithm successfully recovers both the smooth temporal profile of the NCRFs and the spatial extent and location of the active sources, and provides estimates that closely resemble the ground truth. The two-stage localized TRFs, however, fail to recover the true extent of the sources due to the destructive propagation of biases: MNE-boosting estimates are spatially dispersed while Champagne-boosting estimates are overly sparse. Also, the poor signal-to-noise ratio caused the estimates to exhibit spurious peaks in the anterior temporal and inferior frontal lobes: the prominence of these spurious peaks in the Champagne-boosting estimates results in fully overshadowing the true sources. In addition, the two-stage localized TRFs are rescaled using the standard deviation of the sources before plotting. This rescaling, combined with the poor signal-to-noise ratio, leads to the large scaling discrepancy between the estimates and the ground truth. It is worth noting that despite the fact that the Champ-Lasso algorithm is unaware of the true dipole orientations, the resulting NCRF orientations closely match the normal directions of the patches (see Fig. 4). The Champ-Lasso also successfully suppresses spurious peaks in the anterior temporal and inferior frontal lobes, demonstrating its robustness to background activity.

**Figure 4:**
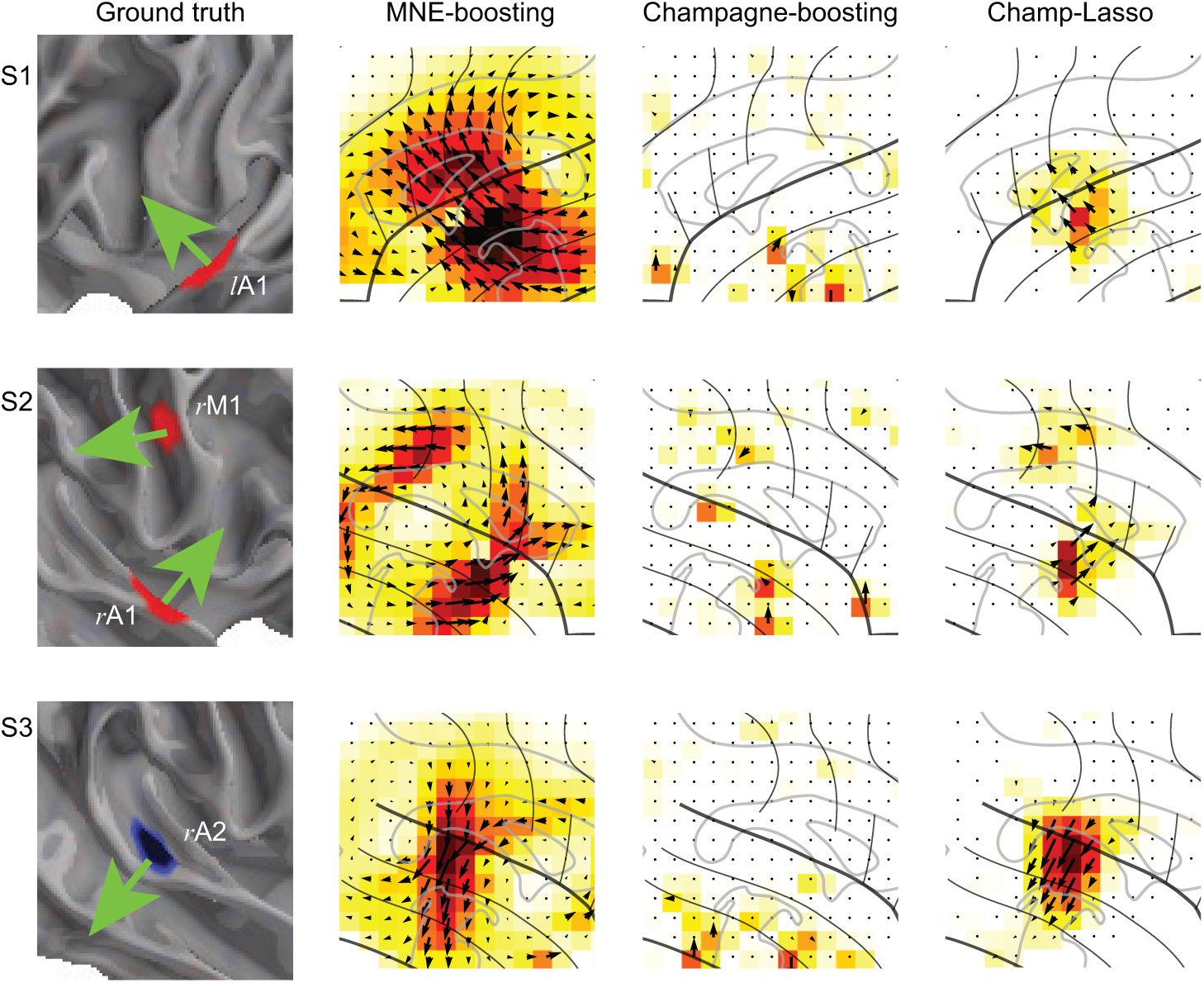
Results for the simulated auditory experiment (continued): Zoomed in views of the active cortical patches (marked as S1, S2 and S2 in Fig. 3A) emphasizing the orientations of the simulated current dipoles (green arrows) alongside the estimated current dipole directions.

Benchmarking metrics described in Section 2.11 are listed in Table 1 and Table 2. Table 1 lists the *dispersion* metric, demonstrating how the different algorithms perform in localizing the neural sources correctly. Table 2 contains the *correlation* measures for the principal orientation and time course along with the selectivity of these principal orientations and time courses across the different simulated regions. While the principal orientations of Champ-Lasso and MNE-boosting are similarly correlated with the true orientation, the selectivity of this orientation in MNE-boosting is inferior to Champ-Lasso. Champagne-boosting exhibits the poorest performance overall. Unlike the other methods, the Champ-Lasso principal time courses consistently show high correlation with the ground truth.

**Table 2:** Comparison with respect to the reconstruction metrics: Pearson correlation coefficients between the estimated *principal orientation* and *principal time course* and the ground truth, as well as their selectivity (higher is better) for different cortical patches (*l*A1, *r*A1, *r*M and *r*A2 as in Fig. 4). The bold numerical values indicate the best performance among the different estimation methodologies in each category.

| | | $lA1$ | $rA1$ | $rM$ | $rA2$ |
| --- | --- | --- | --- | --- | --- |
| Principal Orientation Correlation | Champ-Lasso | **0.991** | **0.992** | **0.923** | **0.996** |
|  | MNE-boosting | 0.978 | 0.989 | **0.923** | 0.995 |
|  | Champagne-boosting | 0.124 | 0.207 | -0.395 | 0.039 |
| Principal Time Course Correlation | Champ-Lasso | 0.968 | **0.953** | **0.972** | **0.958** |
|  | MNE-boosting | 0.959 | 0.856 | 0.741 | 0.918 |
|  | Champagne-boosting | **0.994** | 0.131 | 0.114 | 0.015 |
| Selectivity | Champ-Lasso | **0.999** | **0.977** | **0.932** | 0.984 |
|  | MNE-boosting | 0.977 | 0.794 | 0.469 | 0.845 |
|  | Champagne-boosting | 0.993 | 0.891 | 0.904 | **1.000** |

**Table 1:**
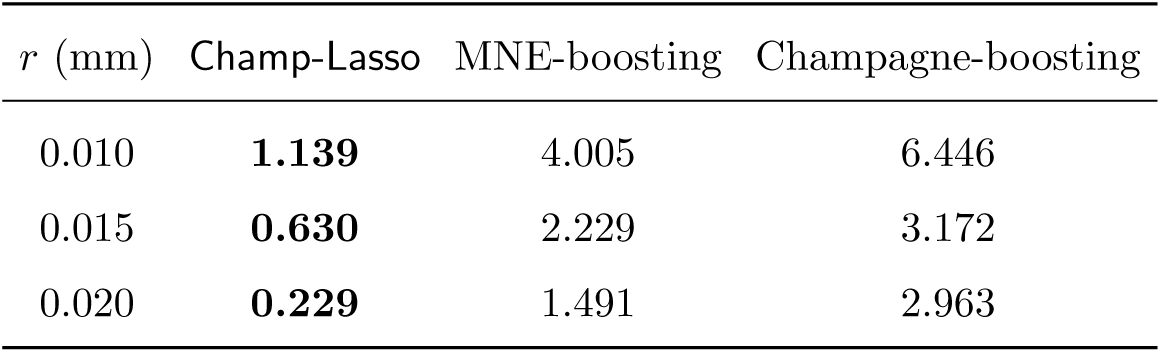
Comparison with respect to the dispersion metric, defined as the ratio of the total NCRF power outside and inside of spheres of radius *r* (for *r* = 10, 15, 20 mm) around the center of mass of the simulated cortical patches (lower is better). The bold numerical values indicate the best performance among the different estimation methodologies for each sphere radius.

| $r$ (mm) | Champ-Lasso | MNE-boosting | Champagne-boosting |
| --- | --- | --- | --- |
| 0.010 | **1.139** | 4.005 | 6.446 |
| 0.015 | **0.630** | 2.229 | 3.172 |
| 0.020 | **0.229** | 1.491 | 2.963 |

### 3.2. Application to Experimentally Recorded MEG Data

#### 3.2.1. Analysis of the Acoustic Envelope NCRFs

Fig. 5A depicts the group average of estimated NCRFs, corresponding to the acoustic envelope, masked by a significance level of *p* = 0.05. The time traces show the magnitude of the average NCRFs (gray segments are statistically insignificant) and the anatomical plots show the spatial NCRF profile at selected temporal peaks, with direction of the vectors projected onto the lateral plane. Fig. 5B shows the temporal profiles (masked at a significance level of *p* = 0.05) of 6 selected NCRFs exhibiting peak spatial activity (collapsed across time). The colored dots on the anatomical plots show the locations of these NCRFs, with matching colors to those of the traces. The traces are grouped by hemisphere and dorsoventrally ordered. The left and right NCRFs in the motor areas are referred to as *l*M_env_ and *r*M_env_, respectively. The left and right auditory NCRF pairs are labeled as *l*A1_env_, *l*A2_env_ and *r*A1_env_, *r*A2_env_, respectively. The NCRFs corresponding to the acoustic envelope in Fig. 5A exhibit two prominent temporal peaks: an early peak at around 30–35 ms, bilaterally centered over the auditory cortex (AC), and a later peak at around 100 ms, dorsal to the first peak and stronger in the right hemisphere. The latter is evident from comparing the left temporal profiles *l*A1_env_ and *l*A2_env_, with their right hemisphere counterparts *r*A1_env_ and *r*A2_env_. Note that the orientations of the NCRFs at the second peak (blue arrow, bottom panel of Fig. 5A) are nearly the opposite of those at the first peak (red arrow, bottom panel of Fig. 5A), which accounts for the negative polarity of the M100 peak with respect to M50 in standard TRF analysis. Furthermore, after the appearance of the first peak (∼ 35 ms, auditory traces *l*A1_env_, *l*A2_env_, *r*A1_env_, and *r*A2_env_) at the AC, the activity appears to gradually shift towards the primary motor cortex (PMC) in both hemispheres (∼ 50 ms, motor traces *l*M_env_ and *r*M_env_). Additionally, the NCRFs show small bilateral late auditory components at around ∼ 250–350 ms.

**Figure 5:**
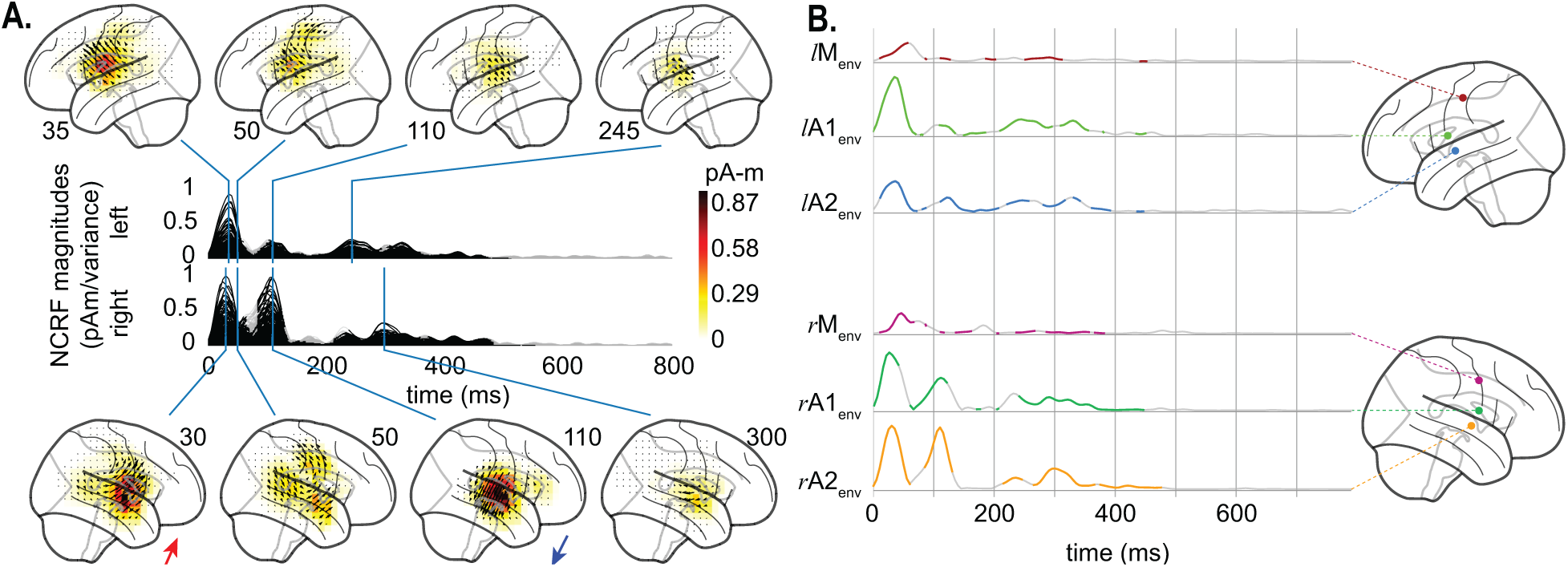
Estimated NCRFs for the acoustic envelope feature variable. A. The anatomical plots show the group-level average NCRFs projected onto the lateral plane (top and bottom panels) corresponding to selected visually salient peaks in the temporal profiles (middle panels). The top and bottom portions of the subplot pertain to left and right hemisphere, respectively. Numerical labels of each anatomical subplot indicates the corresponding time lag in ms. B. The time traces show the temporal profile of 6 selected NCRFs exhibiting peak spatial activity (collapsed across time), grouped by hemisphere and dorsoventrally ordered. The locations of the selected NCRFs are shown on the anatomical plots, with colors matching the time traces and linked by dashed lines. The gray portions of the traces in both subplots indicate statistically insignificant NCRFs at the group level (significance level of 5%). Note that the last 200 ms segments of the temporal profiles are cropped, as they did not correspond to any significant components at the group level. The prominent NCRFs consist of a bilateral auditory component at ∼ 30−35 ms, a bilateral motor component at ∼ 50 ms, and an auditory component at ∼ 110 ms (stronger in right hemisphere and with nearly opposing polarity with respect to the earlier auditory component, indicated by the two colored arrows pointing to the average direction of the NCRFs in subplot A). See Supplementary Movie 01 for a detailed animation showing how the acoustic envelope NCRF components change as function of time lags.

#### 3.2.2 Analysis of the Word Frequency NCRFs

Fig. 6A shows the NCRFs for the word frequency feature variable, in the same format as in Fig. 5. Fig. 6B shows the temporal profiles (masked at a significance level of *p* = 0.05) of 4 selected NCRFs exhibiting peak spatial activity (collapsed across time) also in the same format as in Fig. 5. These include a left auditory (*l*A_wf_), a left frontal (*l*F_wf_), a left inferior temporal (IT) (*l*IT_wf_), and a right auditory (*r*A_wf_) NCRF. The significant NCRF components manifest predominantly in the left hemisphere (Fig. 6A). The earliest peak in the left AC occurs at around 50 ms, followed by a much stronger peak at around 150 ms, slightly posterior to the former (see *l*A_wf_ in Fig. 6B). The earlier peak also has contributions from the inferior temporal gyrus, as indicated by *l*IT_wf_. In addition, the left frontal cortex exhibits weak activity at around 150 ms (see *l*F_wf_ in Fig. 6B). A weak but localized peak centered over the left superior temporal sulcus (STS) is visible at around 240 ms. The only significant component in the right hemisphere occurs around the same time. Finally, the late NCRF components (at around 500–600 ms) mostly originate from the left AC and STS, with weak contributions from the right frontal cortex.

**Figure 6:**
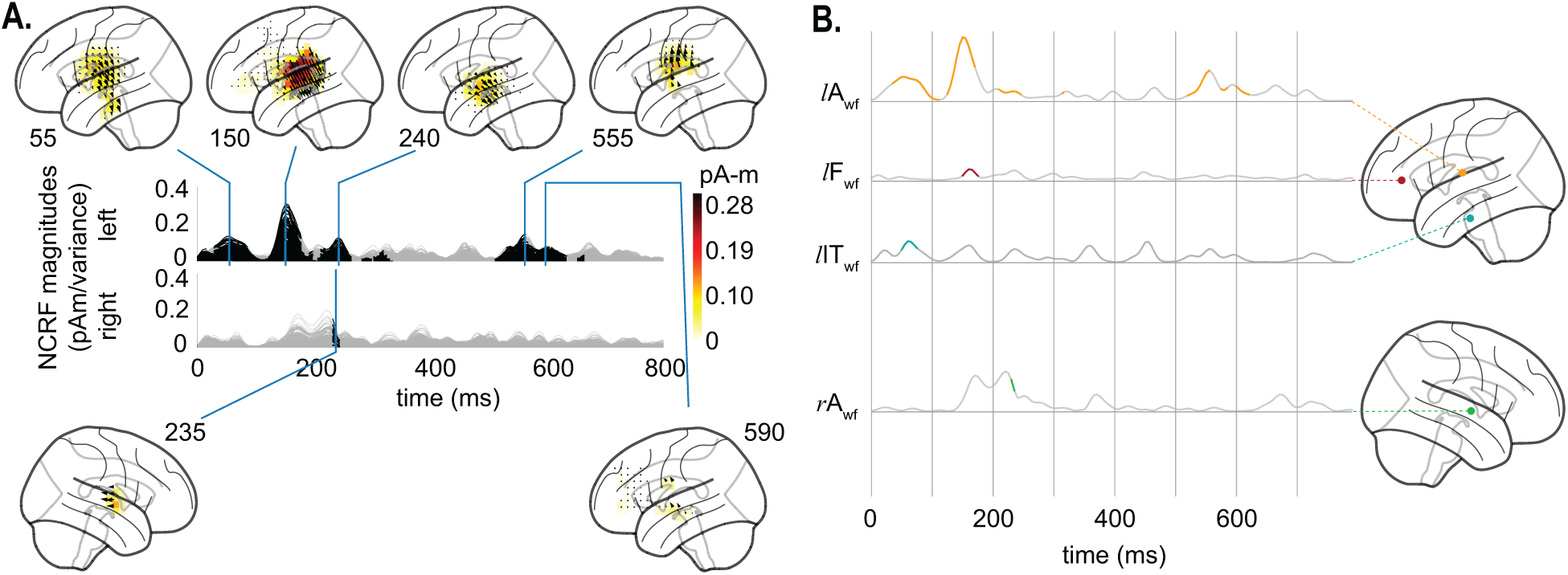
Estimated NCRFs for the word frequency feature variable. A. The anatomical plots show the group-level average NCRFs projected onto the lateral plane (top and bottom panels) corresponding to selected visually salient peaks in the temporal profiles (middle panels). The top and bottom portions of the subplot pertain to left and right hemisphere, respectively. Numerical labels of each anatomical subplot indicates the corresponding time lag in ms. B. The time traces show the temporal profile of 4 selected NCRFs exhibiting peak spatial activity (collapsed across time), grouped by hemisphere and and dorsoventrally ordered. The locations of the selected NCRFs are shown on the anatomical plots, with colors matching the time traces and linked by dashed lines. The anatomical plots show the locations of the selected NCRFs with colors matching the time traces. The gray portions of the traces in both subplots indicate statistically insignificant NCRFs at the group level (significance level of 5%). The prominent NCRFs manifest in the left hemisphere, dominated by an auditory component at ∼ 150 ms. See Supplementary Movie 02 for a detailed animation showing how the word frequency NCRF components change as function of time lags.

#### 3.2.3. Analysis of the Semantic Composition NCRFs

The estimated NCRFs corresponding to the semantic composition feature variable are shown in Fig. 7A, along with 5 representative NCRFs in Fig. 7B. These include two left auditory (*l*A1_sc_ and *l*A2_sc_), two right frontal (*r*F1_sc_ and *r*F2_sc_), and one right middle temporal (*r*MT_sc_) NCRF. The main NCRF components in the left AC peak at around 155 ms and 475 ms, with the earlier peak being ventral to the later one (see *l*A1_sc_ and *l*A2_sc_ in Fig. 7B). The significant right hemispheric NCRFs are temporally concentrated between 155 to 210 ms, and appear superior to those in the left hemisphere, involving inferior frontal gyrus (IFG). Strikingly, these NCRFs in the right hemisphere seem to move in the anterosuperior direction until around 185 ms, at which point the right hemisphere exhibits strong frontal activity (Fig. 7A). The NCRFs return to their initial location afterwards at around 210 ms. This sequence of spatiotemporal changes is also evident in the sequence of temporal peaks in Fig. 7B, given by *r*MT_sc_ → *r*F2_sc_ → *r*F1_sc_ → *r*F2_sc_.

**Figure 7:**
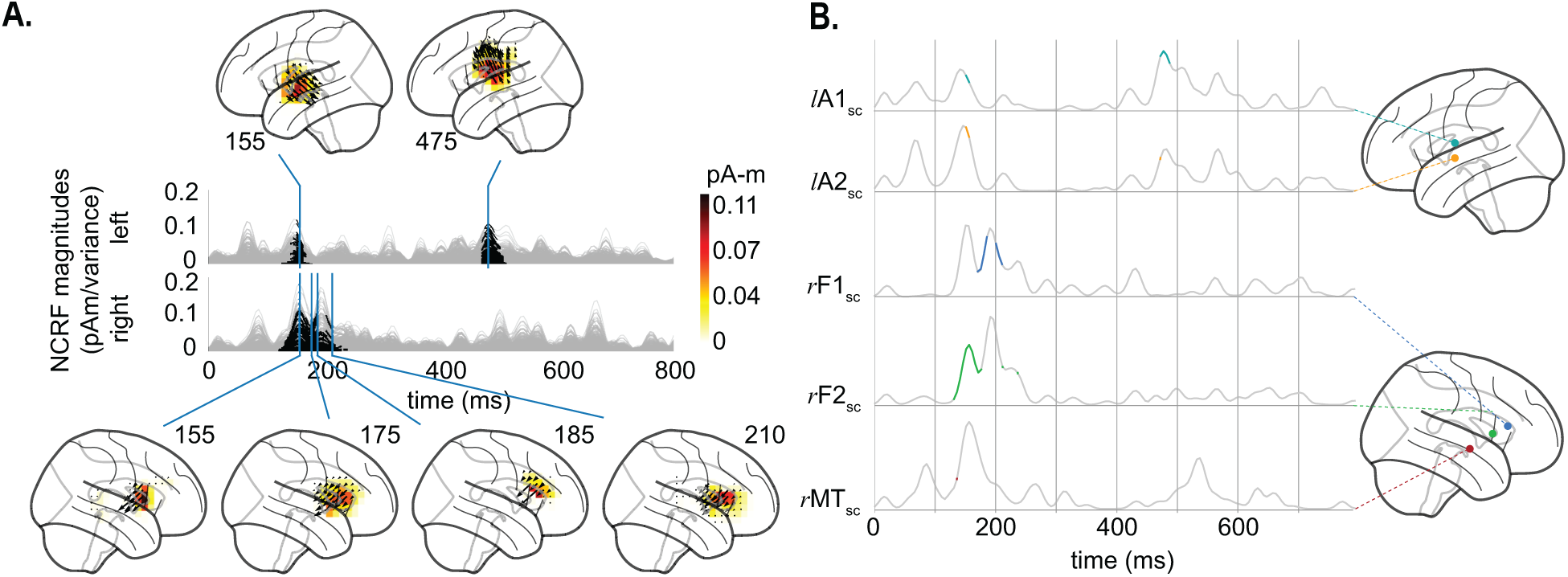
Estimated NCRFs for the semantic composition feature variable. A. The anatomical plots show the group-level average NCRFs projected onto the lateral plane (top and bottom panels) corresponding to selected visually salient peaks in the temporal profiles (middle panels). The top and bottom portions of the subplot pertain to left and right hemisphere, respectively. Numerical labels of each anatomical subplot indicates the corresponding time lag in ms. B. The time traces show the temporal profile of 5 selected NCRFs exhibiting peak spatial activity (collapsed across time), grouped by hemisphere and dorsoventrally ordered. The locations of the selected NCRFs are shown on the anatomical plots, with colors matching the time traces and linked by dashed lines. The gray portions of the traces in both subplots indicate statistically insignificant NCRFs at the group level (significance level of 5%). The prominent NCRF components consist of a bilateral auditory component at ∼ 155 ms, a right auditory-frontal component from ∼ 180 ms to ∼ 210 ms, and a left auditory late component at ∼ 475 ms. The sequence of peaks given by *r*MT_sc_ → *r*F2_sc_ → *r*F1_sc_ → *r*F2_sc_ show the back and forth movement of the NCRFs from the auditory to frontal cortices. See Supplementary Movie 03 for a detailed animation showing how the semantic composition NCRF components change as function of time lags.

## 4. Discussion and Concluding Remarks

Characterizing the dynamics of cortical activity from noninvasive neuroimaging data allows us to probe the underlying mechanisms of sensory processing at high spatiotemporal resolutions. In this work, we demonstrated a framework for direct estimation of such cortical dynamics in response to various features of continuous auditory stimuli from the MEG response. To this end, we developed a fast inverse solution under a Bayesian estimation setting, the Champ-Lasso algorithm, for inferring the Neuro-Current Response Functions (as spatiotemporal models of cortical processing) in a robust and scalable fashion.

One of the key features of the Champ-Lasso algorithm is the ability to simultaneously estimate cortical source covariances in a data-driven fashion (as opposed to relying on data-agnostic depth-weighting procedures) and finding the NCRF model parameters. The interplay between the two as well as incorporating the structural properties of the NCRFs into the model, taking advantage of the Bayesian nature of the estimation framework, ultimately leads to spatially focal NCRFs, with smooth temporal profiles. In other words, the NCRF and source covariance estimation procedures work in tandem to best explain the observed MEG data while minimizing the spatial leakage and capturing the smoothness of the temporal responses. In contrast, previously existing methodologies result in estimates that are spatially broad, which then require post-hoc clustering procedures to meaningfully summarize the underlying spatiotemporal cortical dynamics. These serialized procedures in turn introduce biases to the estimates, and hinder meaningful statistical interpretation of the results.

To demonstrate the utility of our proposed framework, we estimated NCRFs corresponding to several feature variables of speech, reflecting different levels of cognitive processing and comprehension from MEG. The data analyzed in this paper were analyzed by an earlier method in (Brodbeck et al., 2018b), where a two-stage procedure was utilized to probe the cortical processing of speech: the MEG data was first cortically localized using an MNE inverse solver, followed by estimating individual temporal response functions for each source. In order to summarize the resulting estimates in a meaningful fashion, yet another processing step was necessary to disentangle the different spatially dispersed and highly overlapping cortical sources. Our results corroborate those obtained in (Brodbeck et al., 2018b), while obviating the need for any such post-processing, by providing a one-step estimation procedure with the substantial benefit of greatly improved spatial resolution. In addition, the three-dimensional nature of the NCRFs in our framework allows the segregation of different spatial activation patterns that are temporally overlapping. For example, the bilateral activity components in the primary motor cortex in response to the acoustic envelope are automatically clearly distinguishable from the early activation in the auditory cortex, without the need for any post-hoc processing. To ease the visual comparison, Fig. 8 compares the estimated NCRF distributions (transparent cortex) to those of Brodbeck et al. (2018b) (inflated cortical surface), at several time points for each of the three stimulus features.

**Figure 8:**
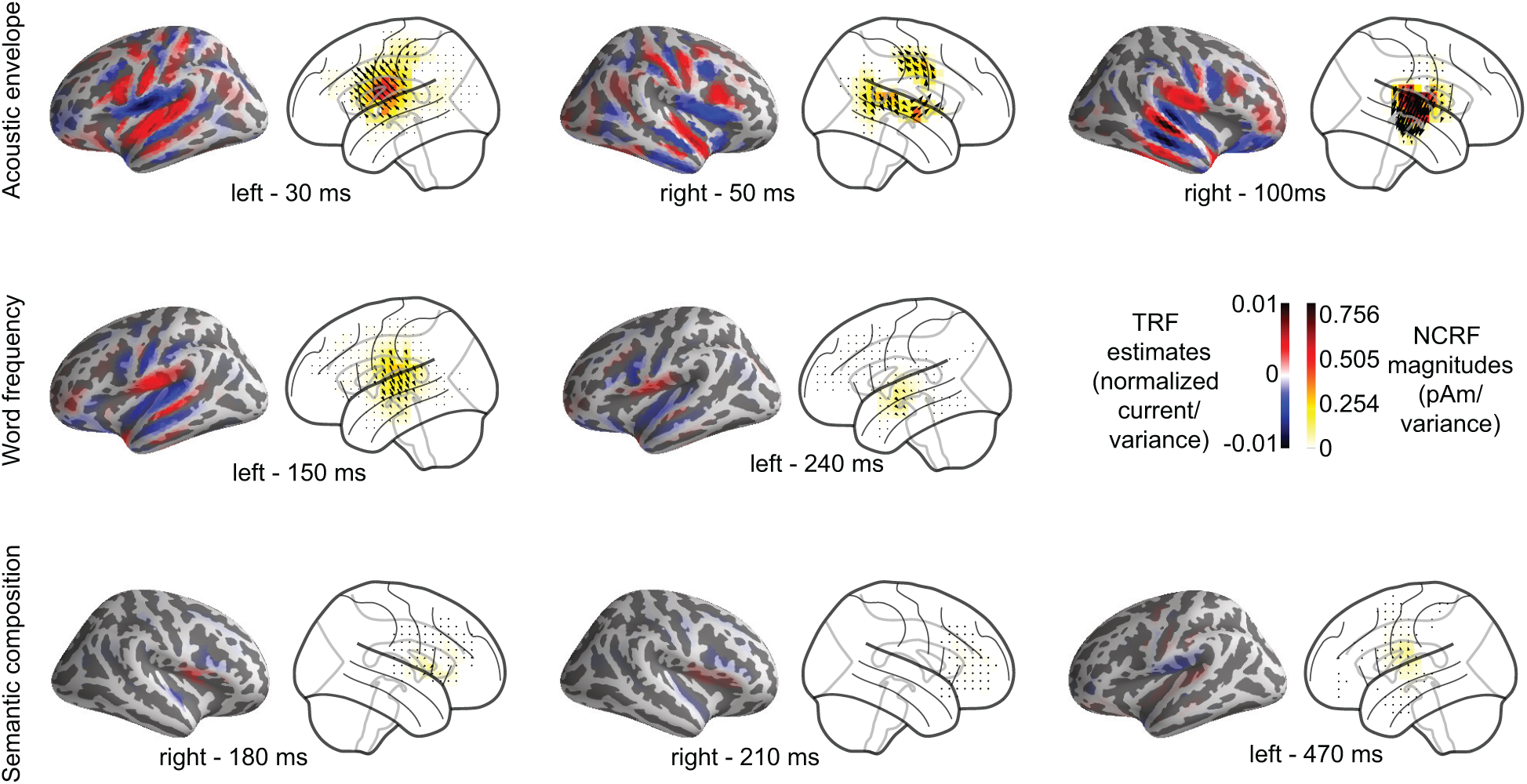
Comparison of cortical spread of estimated NCRFs against MNE-boosting TRFs. The pairwise anatomical plots show the group-level average MNE-boosting TRFs from Brodbeck et al. (2018b) (left) alongside the group-level average NCRFs projected onto the lateral plane (right) for a few selected visually salient peaks in the temporal profiles, corresponding to acoustic envelope (top), word frequency (middle) and semantic composition (bottom) feature variables.

Our results also support other neuroimaging evidence for the hierarchical model of speech processing, involving not only the temporal lobe, but also the motor and frontal cortices (Scott and Johnsrude, 2003; Hickok and Poeppel, 2004; Davis and Johnsrude, 2007; Okada et al., 2010; Peelle et al., 2010; de Heer et al., 2017). To probe the functional organization of this hierarchy, we estimated NCRFs corresponding to features extracted from speech at the acoustic, lexical and semantic levels and found distinct patterns of cortical processing at high spatiotemporal resolutions. Our results indeed imply that while the acoustic and lexical features are processed primarily within the temporal and motor cortical regions (Fadiga et al., 2002; Wilson et al., 2004; Pulvermüller et al., 2006; Crinion et al., 2003; Dewitt and Rauschecker, 2012; Mesgarani et al., 2014; Hullett et al., 2016), phrase-level processing, assessed here using the semantic composition variable, is carried out through the involvement of the frontal cortex (Kaas and Hackett, 1999; Hickok and Poeppel, 2007; Rauschecker and Scott, 2009).

Another advantage of our proposed methodology is mitigating the dependence of the solution on the precise geometry of the underlying cortical source models. In conventional neuromagnetic source imaging, individual structural MR images are utilized in the construction of source space models, particularly for retrieving the cortical surface segmentation. The normal direction to the so-called cortical patches in these models is key in determining the lead-field matrix, which are often referred to as orientation-constrained source models. However, in many available neuroimaging datasets (including the one analyzed in this work), MR images are not available, relying only on an average head model, instead of one informed by the subject-specific cortical geometry. In order to mitigate the need for such information, we utilized a free-orientation volumetric source space in our estimation framework. While this makes the underlying optimization problem more involved and computationally intensive, it adds more than a compensatory amount of flexibility to the underlying models and allows them to recover missing information regarding the cortical source space geometry. To this end, we used rotationally invariant sparsity-inducing priors to regularize the spatiotemporal distribution of the NCRFs. Together with the aforementioned data-driven source covariance adaptation, this regularization scheme results in consistent source orientation estimates and provides a degree of immunity to unwanted side-effects of error-prone coordinate-frame rotations. To confirm these theoretical expectations, we validated this feature of our framework using simulation studies with known ground truth. In light of the above, posing NCRF estimation over an orientation-free volumetric source space can also be thought of as unifying the virtues of distributed source imaging and single dipole fitting: we aim at estimating both the orientations and magnitudes of spatially sparse dipole currents within the head volume that can best linearly predict the MEG responses to continuous stimuli.

This flexibility encourages applications of Champ-Lasso algorithm beyond MEG, for example, to EEG or simultaneous M/EEG recordings. In theory, any source localization method is equally applicable to all such scenarios (albeit with varying performance, due to the intrinsic differences between MEG and EEG), once the lead-field matrix is computed precisely. The main challenge is thus the placement of the current dipoles over the cortical mantle and correctly inferring the orientation of the dipoles from the structural MR scans. Unfortunately, a large majority of EEG experiments do not contain structural MR scans, eliminating the possibility of precise source-space analysis. Our analysis pipeline could be particularly useful for these scenarios, as the particular formulation aims to eliminate this strict requirement on dipole placements by making the solution robust against the unavailability of the precise geometry of the cortical mantle. The favorable performance of the Champ-Lasso algorithm in application to MEG data gives promise of its utility in application to EEG or simultaneous M/EMEG recordings, which would still need to be verified in future studies.

To facilitate such verification as well as usage by the broader systems neuroscience community a Python implementation of the Champ-Lasso algorithm is archived on the open source repository Github (Das, 2019). The current implementation of our algorithm uses the aforementioned regularization scheme to recover temporally smooth and spatially sparse NCRFs. Due to the plug-and-play nature of the proposed Bayesian estimation framework, one can easily utilize other relevant regularization schemes to promote spatial smoothness or incorporate spectro-temporal prior information, by just modifying the penalty term.

## Appendix A. Marginalization

To obtain the marginal distribution of Eq. (9) from the joint distribution of Eq. (8), one needs to integrate out **J** from the latter. Alternatively, thanks to the Gaussian assumption in Eq. (5) and Eq. (7), the marginalization can be carried out as follows. We start from the probabilistic generative model:

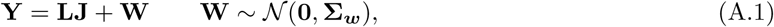

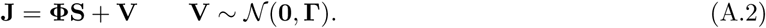

Substituting the expression for **J** from Eq. (A.2) in Eq. (A.1), we arrive at:

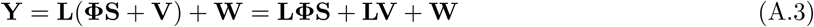

Using the independence of **V** and **W**, the distribution of the stimulus independent part can be derived as **LV** + **W** ∼ 𝒩 (**0, Σ**_***w***_ + **LΓL**^T^). From here, the marginal distribution of **Y** can be written as given by Eq. (9).

## Appendix B. Details of the Regularization Scheme

In this appendix, we provide more details on the regularization scheme used for NCRF estimation. Recall that the NCRF matrix estimation amounts to the following maximum likelihood problem:

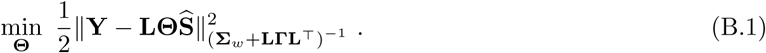

given a particular choice of **Γ**. With this choice, one can find the gradient of the objective as:

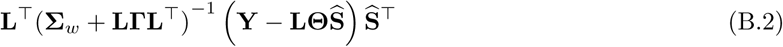

and thus can attempt to solve the maximum likelihood problem using gradient descent techniques. The following observations on the gradient, however, show that the problem is ill-conditioned:

1. The left multiplier of **Θ**, i.e., **L**^T^(**Σ***w* + **LΓL**^T^)^−1^**L** is singular.
2. The right multiplier of **Θ**, i.e, 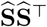, which is the empirical stimulus correlation matrix is likely to be rank-deficient for naturalistic stimuli (Crosse et al., 2016).

Therefore, a direct attempt at solving the problem via the gradient descent results in estimates of **Θ** with high variability. In estimation theory, such ill-conditioning is handled by introducing a bias to the estimator, which contains a priori information about the problem, in order to reduce the estimation variance. In addition, the NCRF model typically has many more free parameters than the observed data points, and without introducing prior information, the estimation problem is prone to over-fitting.

The prior information is often incorporated in the form of regularization. A commonly used regularization scheme in this context is the Tikhonov regularization and its variants for promoting smoothness (Lalor et al., 2006). Other estimation schemes such as boosting and *ℓ*_1_-regularization promote sparse solutions (David et al., 2007; Akram et al., 2017). Here, we introduce a structured regularization by penalizing a specific mixed-norm of the NCRF matrix to recover spatio-temporally sparse solutions over the Gabor coefficients:

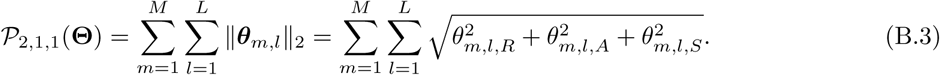

In words, for each current dipole location, we penalize the vector-valued response function by sum of the magnitude of its corresponding Gabor coefficients.

Note that the *ℓ*_1_-regularization in this case, i.e., 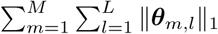, is not compatible with the expected cortical distribution of the NCRFs. Since the *ℓ*_1_-norm is separable with respect to the three 3 coordinates of ***θ***_*m,l*_, it tends to select a sparse subset of the 3D coordinates, rendering the recovered NCRF components parallel to the coordinate axes. In contrast, the proposed penalty aims to select the NCRF components as a single entity by penalizing the vector magnitudes at each lag. Indeed, if the current dipoles are constrained to be normal to the cortical patches in the NCRF formulation, the proposed penalty coincides with *ℓ*_1_-regularization.

Another advantage of this mixed-norm penalty is its rotational invariance when working with 3D vector-valued response functions. Suppose the coordinate system is rotated by an orthogonal matrix ***U*** ∈ ℝ^3×3^. Then, the lead-field and NCRF matrices are transformed by: **L** → **L*Ũ***^T^ =: **L**′, **Θ** → ***Ũ* Θ** =: **Θ**′ where ***Ũ*** = ***I***_*M*_ ⊗ ***U***. Then, **L**′**Θ**′ = **L*Ũ*** ^T^***Ũ* Θ** = **LΘ** and

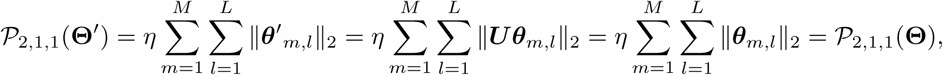

which implies the aforementioned rotational invariance. As a result, the solutions are not dependent on any particular choice of coordinate system. Also, since the penalty does not prefer specific source orientations, it makes the solution more resilient to co-registration error than other approaches that do not consider the vector-valued nature of the current dipoles or constrain the solutions to be normal to the cortical surface.

## Appendix C. Statistical Testing Procedures

To asses the statistical significance of the estimated NCRF components at the group level and across the source space, they need to be compared against suitable null hypotheses. The fact that the NCRF components are 3D vectors requires technical care in choosing the null hypotheses. Here, we provide two possible null hypotheses and testing methodologies: the Length Test, that only considers the length or magnitude of the NCRFs, and the Vector Test that takes into account both the magnitude and direction of the NCRFs. The corresponding source codes that implement these tests can be found at (Brodbeck, 2017).

### The Length Test

This test aims to assess the statistical significance of the NCRF components by comparing their magnitudes against a baseline ‘null’ NCRF model at the group level. To control for false positives arising from over-fitting, instead of using an all-zero null model of the NCRFs, we aim to learn the null model from the dataset itself. The time-series of the feature variables are split into four equal segments, and these segments are permuted cyclically to yield three ‘misaligned’ feature time-series. Then, for each feature variable, three ‘misaligned’ time-series are constructed by swapping its original time-series with the ‘misaligned’ ones, while keeping the other two feature variables intact. Then, the average NCRF magnitudes estimated from these three ‘misaligned’ time-series are considered as the null model for that feature variable. The NCRF magnitude pairs from the original data and the null model are tested for significance using mass-univariate tests based on related measures t-tests.

To control for multiple comparisons, nonparametric permutation tests (Nichols and Holmes, 2002; Maris and Oostenveld, 2007) based on the threshold-free cluster-enhancement (TFCE) algorithm (Smith and Nichols, 2009) are used. First, at each dipole location and time point, the t-statistic is computed from the difference between the NCRF magnitude pairs. The resulting statistic-map is then processed by the TFCE algorithm, which boosts contiguous regions with high test statistic as compared to isolated ones, based on the assumption that spatial extent of the true sources is typically broader than those generated by noise. To find the distribution of these TFCE values under the null hypothesis, TFCE values are calculated following the same procedure, on 10000 different random permutations of the data. In each permutation, the sign of the NCRF magnitude differences is flipped for a randomly selected set of subjects, without resampling the same set of subjects. Then, at every permutation, the maximum value of the obtained TFCE values is recorded, thereby constructing a non-parametric distribution of the maximum TFCE values under the null hypothesis. The original TFCE values that exceed the (1 − *α*) percentile of the null distribution are considered significant at a level of *α* corrected for multiple comparisons across the sources.

### The Vector Test

This test aims at quantifying the significance of the estimated 3D NCRF components at the group level, based on the one-sample Hotellings *T*^2^ test. In the one-sample Hotellings *T*^2^ test, the population mean of the sample vectors is tested against the null hypothesis of mean zero, i.e. ***µ***_0_ = 0. To control for multiple comparisons, a similar strategy based on nonparametric permutations as in the case of the Length Test is used. At every time lag, the *T*^2^ statistic for each dipole is computed as:

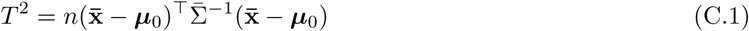

where 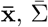 are the population mean and covariance matrix of the vector-valued NCRF components, respectively. The *T*^2^ statistic quantifies the variability of vector-valued samples, akin to the role of the *t*-statistic for 1D samples (Mardia, 1975). The resulting *T*^2^-maps are then processed by the TFCE algorithm. As before, to construct a non-parametric distribution of maximum TFCE values under the null hypothesis, maximum values of the TFCE-processed *T*^2^ maps on 10000 different random permutations of the data are recorded. In each permutation, the vector-valued NCRF components of each subject undergo uniform random rotations in 3D (Miles, 1965). The original TFCE values that exceed the (1 − *α*) percentile of the null distribution are considered significant at a level of *α*, corrected for multiple comparisons across the sources.

Traditionally, response functions are estimated as scalar functions of the data, either over the sensor space or over the source space by orientation-constrained inverse solvers. Considering the directional variability of the NCRF estimates at the group level, however, takes into account the group level anatomical variability that may effect the current dipole orientations. In addition, the Vector Test is less computationally demanding than the Length Test, because it does not require refitting NCRFs for permuted models. In the Results section of the manuscript, we presented the NCRFs masked at a significance level of 5%, based on the Length Test. To demonstrate the difference between these two tests, here we also present the sames results using the Vector Test (Fig C.9).

**Figure C.9:**
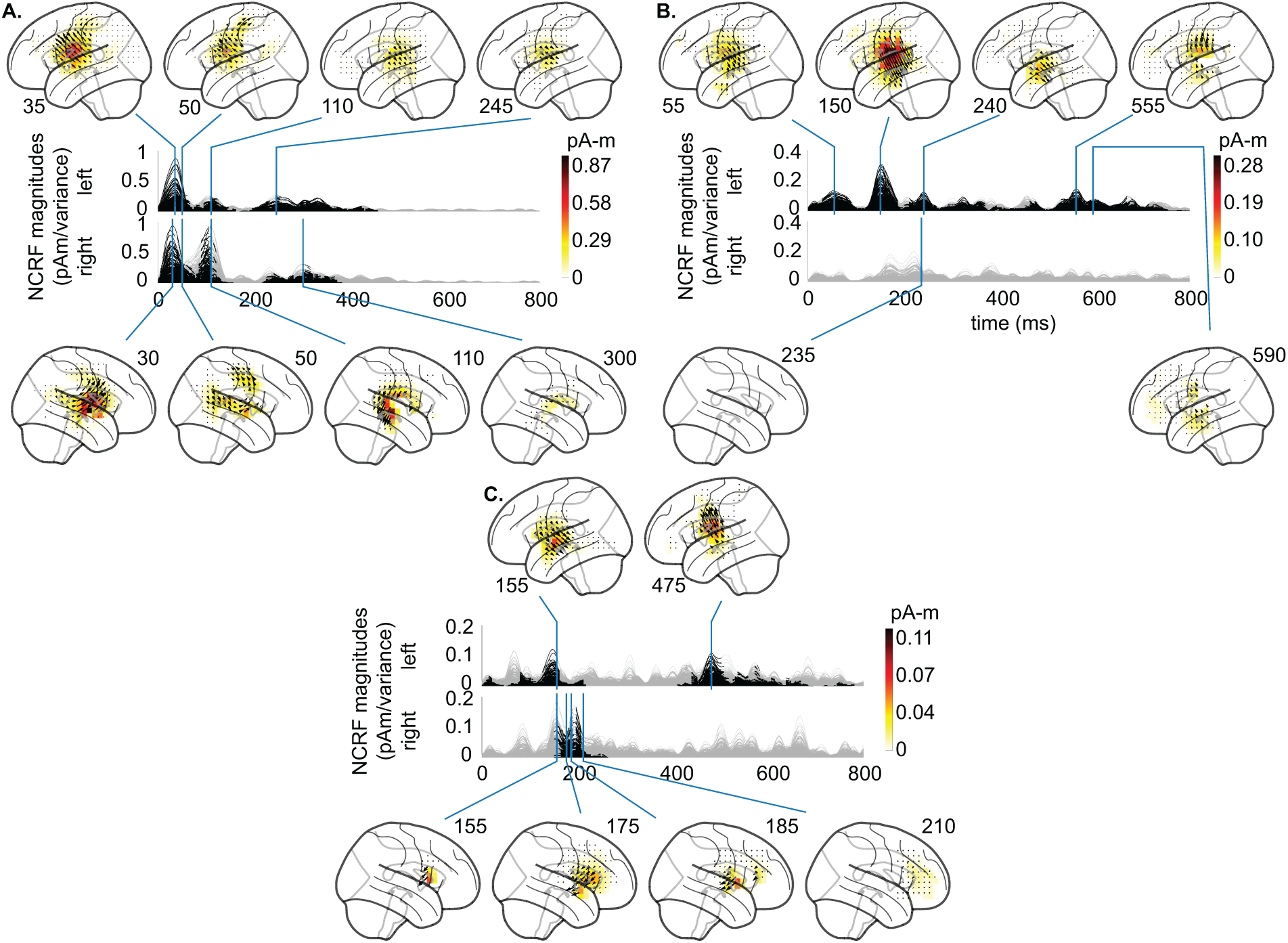
Estimated NCRFs for acoustic envelope (A), word frequency (B), semantic composition (C): The anatomical plots show the group-level average NCRFs projected onto the lateral plane (top and bottom panels) corresponding to selected visually salient peaks in the temporal profiles (middle panels). The top and bottom portions of the subplot pertain to left and right hemisphere, respectively. Numerical labels of each anatomical subplot indicates the corresponding time lag in ms. The gray portions of the traces indicate statistically insignificant NCRFs at the group level (significance level of 5%). The significance levels are computed using the Vector Test, as opposed to the main manuscript where the significance levels are based on the Length Test. The main features of the NCRFs discussed in the Results section are similarly recovered by the Vector Test.

